# Transcriptional control of hypoxic hyphal growth in the fungal pathogen *Candida albicans*

**DOI:** 10.1101/2021.09.02.458602

**Authors:** Manon Henry, Anais Burgain, Faiza Tebbji, Adnane Sellam

**Affiliations:** Montreal Heart Institute, Université de Montréal, Montréal, QC, Canada; Department of Microbiology, Infectious Diseases and Immunology, Faculty of Medicine, Université Laval, Quebec City, QC, Canada; Department of Microbiology, Infectious Diseases and Immunology, Faculty of Medicine, Université de Montréal, Montréal, QC, Canada

**Keywords:** *Candida albicans*, hypoxia, filamentation, transcriptional control, transcriptomics, ChIP-chip, transcription factors

## Abstract

**Background:** The ability of *Candida albicans*, an important human fungal pathogen, to develop filamentous forms is a crucial determinant for host invasion and virulence. Filamentation is triggered by different host environmental cues. Hypoxia, the dominant conditions that *C. albicans* encounters inside the human host, promote filamentation, however, the contributing mechanisms remain poorly characterized.

**Methods:** We performed a quantitative analysis of gene deletion mutants from different collections of protein kinases and transcriptional regulators in *C. albicans* to identify specific modulators of the hypoxic filamentation. We used genome-wide transcriptional profiling (Microarrays) and promoter occupancy (ChIP-chip) to characterize regulons of two transcription factors that were associated with the hypoxic filamentation. Genetic interactions were also used to assess functional relationships among the newly identified modulators of hypoxic filamentation and the well-known *C. albicans* core morphogenetic regulators.

**Results:** Our genetic screen uncovered two transcription factors, Ahr1 and Tye7, that act as prominent regulators of *C. albicans* filamentation specifically under hypoxia. Both *ahr1* and *tye7* mutants exhibited a hyperfilamentous phenotype specifically under an oxygen-depleted environment suggesting that these transcription factors act as negative regulators of hypoxic filamentation. By combining microarray and ChIP-chip data, we have characterized the set of genes that are directly modulated by Ahr1 and Tye7. We found that both Ahr1 and Tye7 modulate a different set of genes and biological processes. Our genetic epistasis analysis supports our genomic finding and suggests that Ahr1 and Tye7 act independently to modulate hyphal growth in response to hypoxia. Furthermore, our genetic interaction experiments uncovered that Ahr1 and Tye7 repress the hypoxic filamentation growth via the Efg1 and Ras1/Cyr1 pathways, respectively.

**Conclusion:** In sum, this investigation represents an informative resource toward the understanding of how hypoxia, the predominant condition inside the host, shapes the invasive filamentous growth of *C. albicans*.

## Introduction

*Candida albicans* is an opportunistic fungus and one of the most common causes of systemic fungal infection in humans with high mortality rates of 50% or greater despite currently available antifungal therapy [1,2]. The ability of this yeast to cause infections is centrally related to different intrinsic virulence attributes such as biofilm formation, adherence to the host epithelia, the yeast-to-hyphae switch, secretion of hydrolytic enzymes (e.g. lipases, phospholipases and proteinases) and the cytolytic peptide Candidalysin [3,4]. Of particular importance, a transition from the yeast morphology to the filamentous form is a key virulence determinant dedicated to the invasion of host tissues and, allowing to escape from phagocytes [5]. Hyphal cells are also characterized by an enhanced adhesiveness to mucosal surfaces and are essential for the compression strength of *C. albicans* biofilms [6,7]. Additionally, as s in *Saccharomyces cerevisiae*, filamentation might allow *C. albicans* to conquer niches where nutrient conditions are not limiting [8,9].

*C. albicans* yeast-to-hyphae transition is controlled by an intertwined regulatory circuit that signals different filamentation stimuli encountered in distinct host microenvironments including pH, serum, N-acetylglucosamine, temperature, nutritional stress, hypoxia and CO_2_ [10]. Hypoxia, the dominant conditions that *C. albicans* encounters inside the human host, promote filamentation, however, the contributing mechanisms remain poorly characterized. In response to both CO_2_ and hypoxia, Ofd1, a prolyl hydroxylase, contribute to the maintenance of hyphae elongation by stabilizing the transcriptional activator of hypha-specific genes, Ume6 [11]. The transcription factors Efg1 and Bcr1 act both as repressors of filamentation under hypoxic environments to sustain the commensal growth of *C. albicans* at temperatures ≤ 35°C that are slightly below the core body temperature (i.e. < 37°C) [12,13]. In opposite to Efg1-Bcr1 regulatory axis, the transcription factor Ace2 promotes filamentation under hypoxia [14] but also in response to the hyphae-promoting growth medium, Spider [15]. Many other regulatory proteins were also essential for the *C. albicans* morphogenesis under hypoxia, however, they were also required to signal in response to other filamentation cues [16–22].

So far, there are no known regulatory circuits that mediate filamentation exclusively in response to hypoxia. This could be explained by the fact that the *C. albicans* morphogenesis regulatory pathways might be evolutionary optimized to integrate different combinations of filamentation cues to effectively promote invasive growth and virulence. In this study, we performed a quantitative analysis of gene deletion mutants from different collections of protein kinases and transcriptional regulators in *C. albicans* to identify specific regulators of the hypoxic filamentation [15,23–25]. Our results recomforted the aforementioned hypothesis as the majority of mutants with filamentation defects under hypoxia were previously shown to exhibit the same phenotype in response to other hyphae-promoting cues. Our work uncovered two transcription factors, Ahr1 and Tye7, that act as prominent regulators of *C. albicans* filamentation specifically under hypoxia. In summary, we used genome-wide transcriptional profiling and promoter occupancy to characterize both Ahr1 and Tye7 regulons associated with the hypoxic filamentation in *C. albicans*. Our data show that both Ahr1 and Tye7 act as negative regulators of the hypoxia-induced filamentation by modulating a different set of genes. Our genetic epistasis analysis supports our genomic finding and suggests that Ahr1 and Tye7 act independently to modulate hyphal growth in response to hypoxia.

## Results and discussion

### Screening of transcriptional and signaling regulatory mutant libraries for filamentous growth defect under hypoxia

To gain insight into regulatory networks that control hyphae formation in response to hypoxia, a compilation of 370 unique mutants of regulatory proteins related to diverse signaling pathways (140 mutants) and transcriptional regulators (230 mutants) from many publicly available libraries were screened (**Table S1**) [23–25]. As sucrose led to enhanced filamentation under hypoxic environments [14,26], our screens were performed using YPS (Yeast-Peptone-Sucrose) medium at 37°C. Under normoxic conditions, the WT strain formed colonies with short invasive and aerial filaments, while under hypoxic conditions those filaments are invasive and, at least, four-times longer (**Figure 1 A-B**). The ability of each mutant to form hyphae under hypoxic conditions was assessed by scoring the filamentation of peripheral regions of colonies. Wild type morphology was scored as 0, reduction of filamentation was scored from -1 to -3 and hyperfilamentation from 1 to 3. After verifying the observed phenotypes, we have confirmed filamentation defect for 50 mutants in both normoxic and hypoxic conditions. Most of those mutants (39 mutant strains) displayed a reduction of filamentous growth compared to their respective parental strains while 8 strains exhibited enhanced filamentous growth (**Figure 1C-E**). A total of three mutant strains (*ahr1, sch9, tye7*) exhibited altered filamentation specifically under a low oxygen environment (**Figure 1F**).

**Figure 1.**
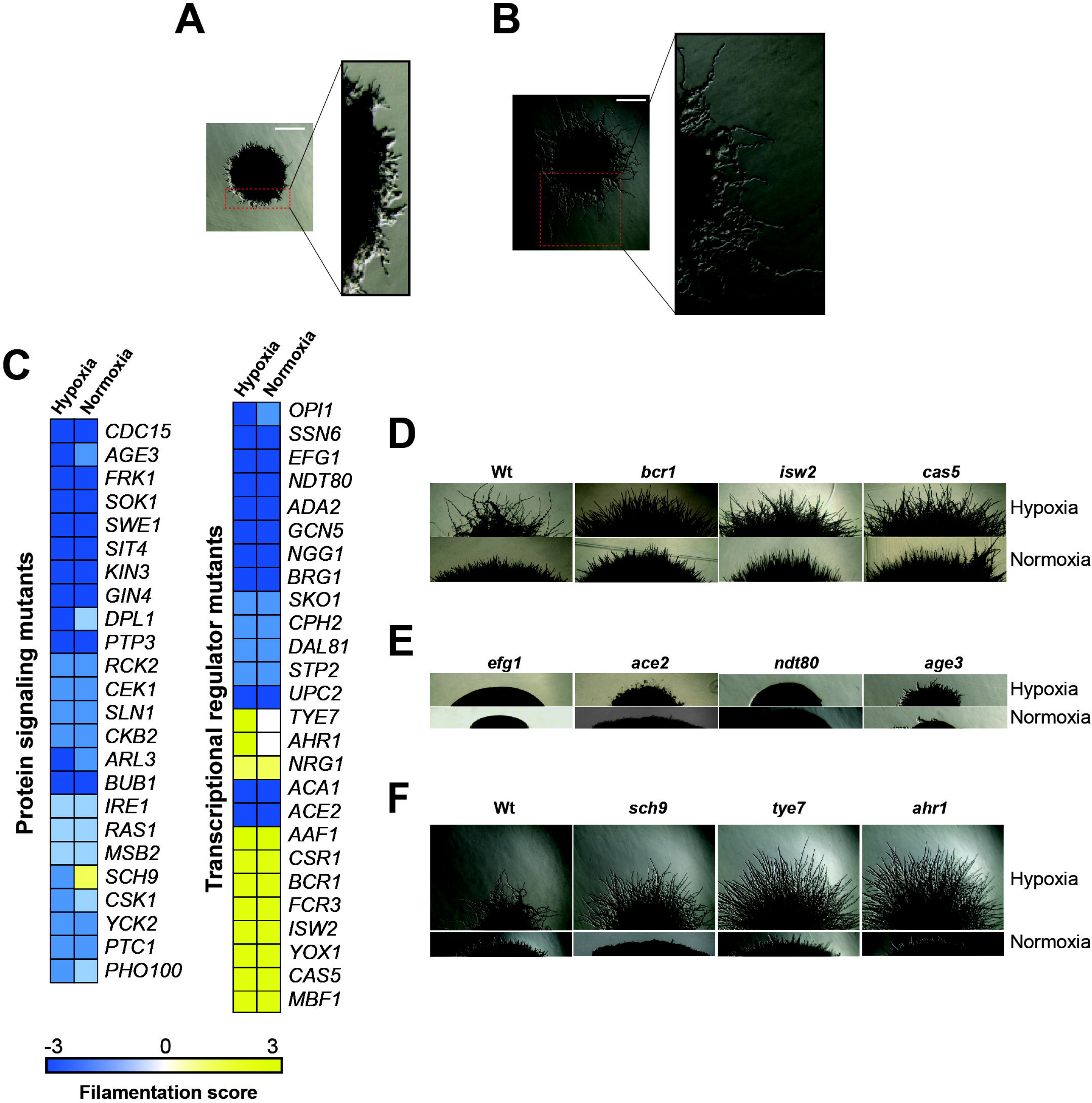
Genetic survey for regulatory proteins required for hypoxic and normoxic filamentation. Morphology of *C. albicans* colony under both normoxia (**A**) and hypoxia (**B**) growing in YPS medium at 37°C. Peripherical regions are magnified and showed short filaments under normoxic conditions (**A**) while under hypoxia these invasive hyphae are ∼ 4 times longer (**B**). (**C**) Filamentation scoring of the 50 identified mutants. WT morphology was scored as 0, reduction of filamentation was scored from -1 to -3 and hyperfilamentation from 1 to 3. (**D-F**) Single-cell– derived colony morphologies representative of hyperfilamentous (**D**) and nonfilamentous (**F**) mutants under either normoxia or hypoxia. (**E**) Mutants with altered filamentation specifically under hypoxia. Bar, 20 µm.

Mutants of transcriptional regulation showing a filamentous growth alteration both under normoxic and hypoxic conditions included many well-characterized hyphal regulators such as Ndt80, Ssn6, Ace2, Upc2, Cph2, Efg1 and the SAGA complex components Ada2 and Gcn5 (**Table S1**). Among mutants with altered hyphal growth, *bcr1, ndt80, upc2, age3* and *ace2* have been already characterized as defectives in filamentation under hypoxic conditions [14,17,27,28]. Filamentation defects of different protein kinase mutants (*ire1, sok1, gin4*) reported in the previous large-scale study by Blankenship *et al*. [24] under normoxic conditions were also confirmed in our screen. Mutants of other signaling components such as phosphatases (*sit4, ptp3*), cell wall sensor (*msb2*), protein acting in the cAMP-mediated (*ras1*) and the two-component signaling pathways (*nik1, sln1*) were significantly defectives in hyphal growth under both normoxic and hypoxic conditions.

### The transcription factors Ahr1 and Tye7 modulate filamentation specifically under hypoxia

Among the transcriptional regulators for which we have defined a role in hyphae formation in this study, are the transcription factors Dal81, Mbf1, Yox1 and Fcr3, and the component of the ADA/SAGA complex Ngg1. We also observed hyphal growth defects in mutants of diverse signaling pathways, whose role in filamentation was not yet demonstrated (Swe1, Cdc15, Bub1, Ptk2, Kin3, Ckb2, Pho100, Csk1 and Dpl1). Mutants displaying enhanced filamentation specifically under low oxygen environments include *ahr1* and *tye7* (**Figure 1F)**. Tye7p and Ahr1p are both transcription factors controlling the expression of glycolytic and adhesin genes, respectively [26,29]. Inactivation of the AGC protein kinase Sch9 led to a hyperfilamentation phenotype comparable to that of *ahr1* and *tye7* under hypoxia, while under normoxia *sch9* cells were unable to differentiate hyphae (**Figure 1F)**. Earlier works have described the hyperfilamentous phenotype of both *tye7* and *sch9* under hypoxia [19,30], however, the exact mechanisms by which Tye7 and Sch9 modulate the hypoxic filamentation remain to be determined. For the current study, we decided to focus on Ahr1 and Tye7 as they represent potent regulators of hyphal growth specifically under hypoxia.

### Transcriptional program driving hypoxic filamentation

First, and prior to assessing the contribution of Ahr1 and Tye7 on transcriptional control of the hypoxia-induced filamentation, we wanted to define transcripts that are differentially regulated when *C. albicans* grow as colonies in a solid YPS medium under hypoxic conditions. Differentially expressed genes were identified by comparing the transcriptional profiles of colonies exposed to low oxygen concentration (1% O_2_) to the transcriptional profiles of colonies growing under normoxic conditions (21% O_2_). Using a statistical significance analysis with an estimated false discovery rate of 5%, in addition to a cut-off of 1.5-fold, we identified 365 hypoxic-responsive genes, including 125 upregulated transcripts and 240 downregulated transcripts (**Table S2**). Upregulated genes were significantly enriched in transcripts related to ribosomal biogenesis and rRNA processing as well as metabolic functions such as amino acid and carboxylic acid biosynthesis (**Figure 2A**). The transcript level of genes related lipid metabolism including ergosterol (*ERG1, ERG5, ERG11, ERG13, DAG7, HMG1, IDI1* and *CYB5*), sphingolipid (*SCS7, DES1, SLD1, FEN1* and *ARV1*), and fatty acids (*FAS1, FAS2, FAD2, FAD3* and *ALK8*) biosynthesis were also significantly induced (**Figure 2A and Table S2**). Repressed transcripts were enriched in many functions, such as ATP synthesis coupled mitochondrial proton transport (*ATP1, ATP2, ATP5, ATP7, ATP14, ATP18 and ATP19*) and nucleosome components (*HTA2, HHF22, HHF1, HTB1, HTB2* and *HTA1*) (**Figure 2B**).

**Figure 2.**
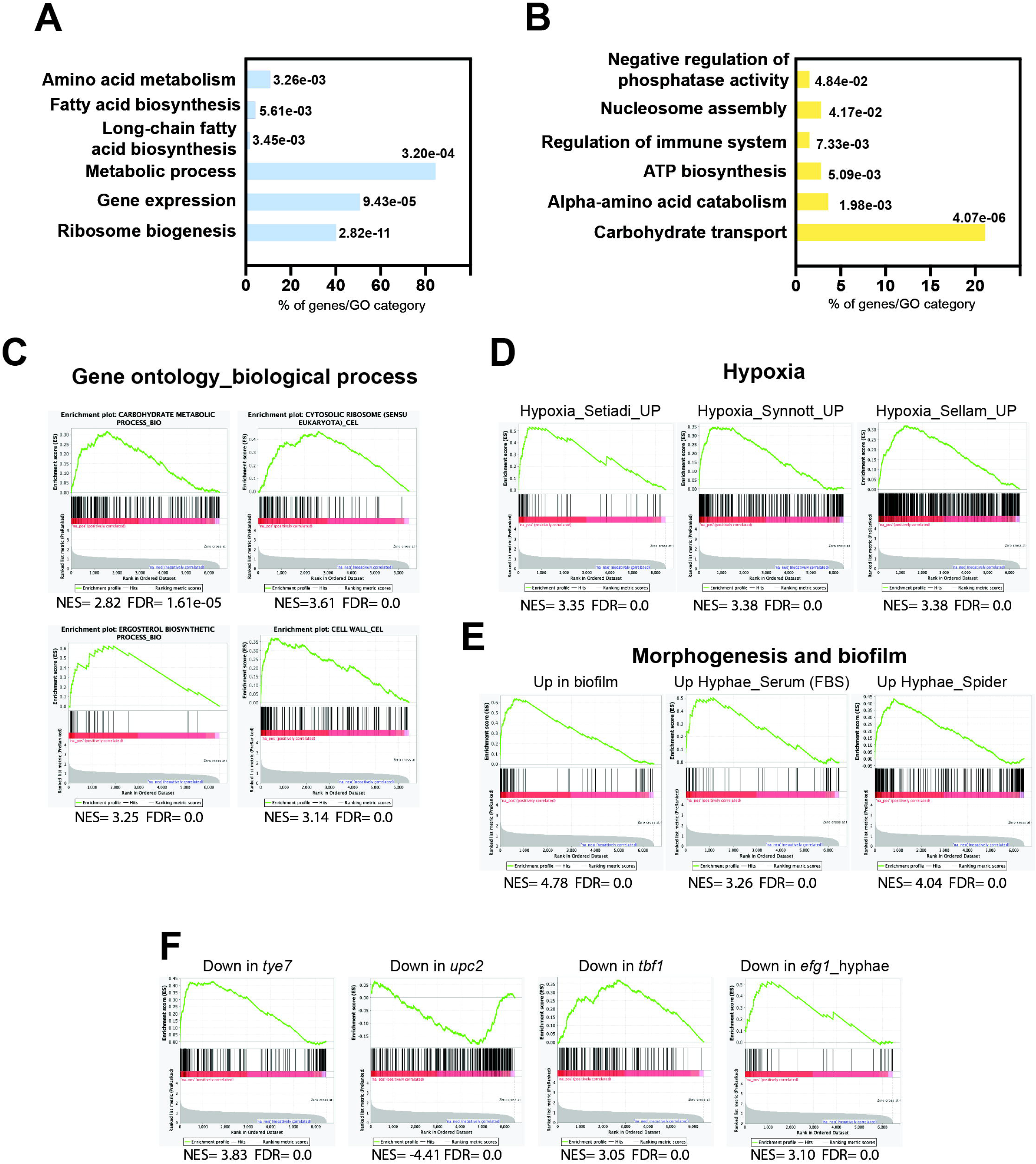
Transcriptional profile driving hypoxic filamentation. Gene ontology analysis of upregulated (**A**) and downregulated (**B**) transcripts of *C. albicans* colonies growing under hypoxia. The *p*-values were calculated using the hypergeometric distribution. (**C-F**) Gene set enrichment analysis of the transcriptional signatures modulated in response to hypoxia. The complete GSEA correlations are listed in **Table S3**. Graphs of GSEA of relevant correlations with different biological functions (**C**), former transcriptomics analysis of *C. albicans* growing under hypoxia (**D**) or under biofilm and hyphal growth states (**E**), and in different mutant’s backgrounds (**F**). NES, normalized enrichment score; FDR (*q*-value): False Discovery Rate.

To further mine the transcriptional program underlying hyphae formation under hypoxia, we have used the Gene Set Enrichment Analysis (GSEA) tool [31,32]. GSEA recapitulated the different biological functions altered under the hypoxic filamentation in addition to underlying the carbohydrate genes enrichment among upregulated transcripts (**Figure 2C; Table S3**). The GSEA analysis of the hypoxic filamentation reflected a complex signature that is similar concurrently to that experienced by yeast cells growing under similar oxygen status [12,28,33], and cells undergoing hyphal growth in response to different cues under normoxia (**Figure 2D-E**). Intriguingly, the enhanced hypoxic filamentation was not accompanied by the activation of the core filamentation genes including the cell wall proteins Hwp1, Als3, Ece1, Ihd1 and Rbt1 [34,35]. This could be explained by the fact that these transcripts are equally expressed both in our microarrays normoxic control and the hypoxic treatment. A significant similarity was also perceived with cells growing as biofilm confirming the hypoxic environment of this sessile growth of *C. albicans* [36] (**Figure 2E**).

GSEA uncovered that the hypoxic filamentation program was similar to that of mutants of different transcription factors including *tye7, upc2* and *tbf1* matching their known role in modulating biological processes that were differentially modulated in our experiment including carbohydrate metabolism, ergosterol biosynthesis and translation, respectively (**Figure 2F** and **Table S3**). Upregulated transcripts displayed a significant correlation with genes requiring the master filamentation regulator Efg1 for their proper activation. This reflects that the enhanced filamentous growth under hypoxia might be driven by Efg1.

While hypoxic filamentation was not supported by the activation of the core filamentation genes, different metabolic processes were differentially modulated reflecting a cellular reprogramming of *C. albicans* metabolism to accommodate the metabolic demand accompanying the enhanced hyphal growth (**Figure 2, Table S2 and S3**). In *S. cerevisiae* and other yeasts, filamentation allows conquering niches where nutrient conditions are not limiting [37]. As hypoxia led to the depletion of many essential metabolites in *C. albicans* [38], the enhanced filamentous growth might consequently represent a nutritional scavenging response as observed in the budding yeast [37]. For instance, oxygen scarcity leads to ergosterol depletion in *C. albicans* [38] which could serve as a cue to promote hyphal growth. In a support of such hypothesis, genetic perturbation of *ERG* genes led to a hypofilamentous phenotype comparable to that observed under hypoxia [39].

### Transcriptional profiling of *tye7* mutant under hypoxia

Among the 43-repressed transcripts in *tye7*, a total of 26 were direct targets of Tye7p as previously reported [26] (**Figure 3A and Table S4**). This includes glycolytic genes (*TPI1, CDC19, GLK1, GPM2, PFK26, PGI1*) and genes related to sugar metabolisms (*HGT8, OSM1, PDC11, PGM2* and *SHA3*) (**Figure 3A and 3C**). This finding recapitulates the well-known role of Tye7 as a major transcriptional regulator of carbohydrate metabolism in *C. albicans* under hypoxia [29,33]. The hyperfilamentation phenotype of *tye7* might be explained by a reduction of glycolytic flux and impairment of carbon metabolism as previously shown [26,30].

**Figure 3.**
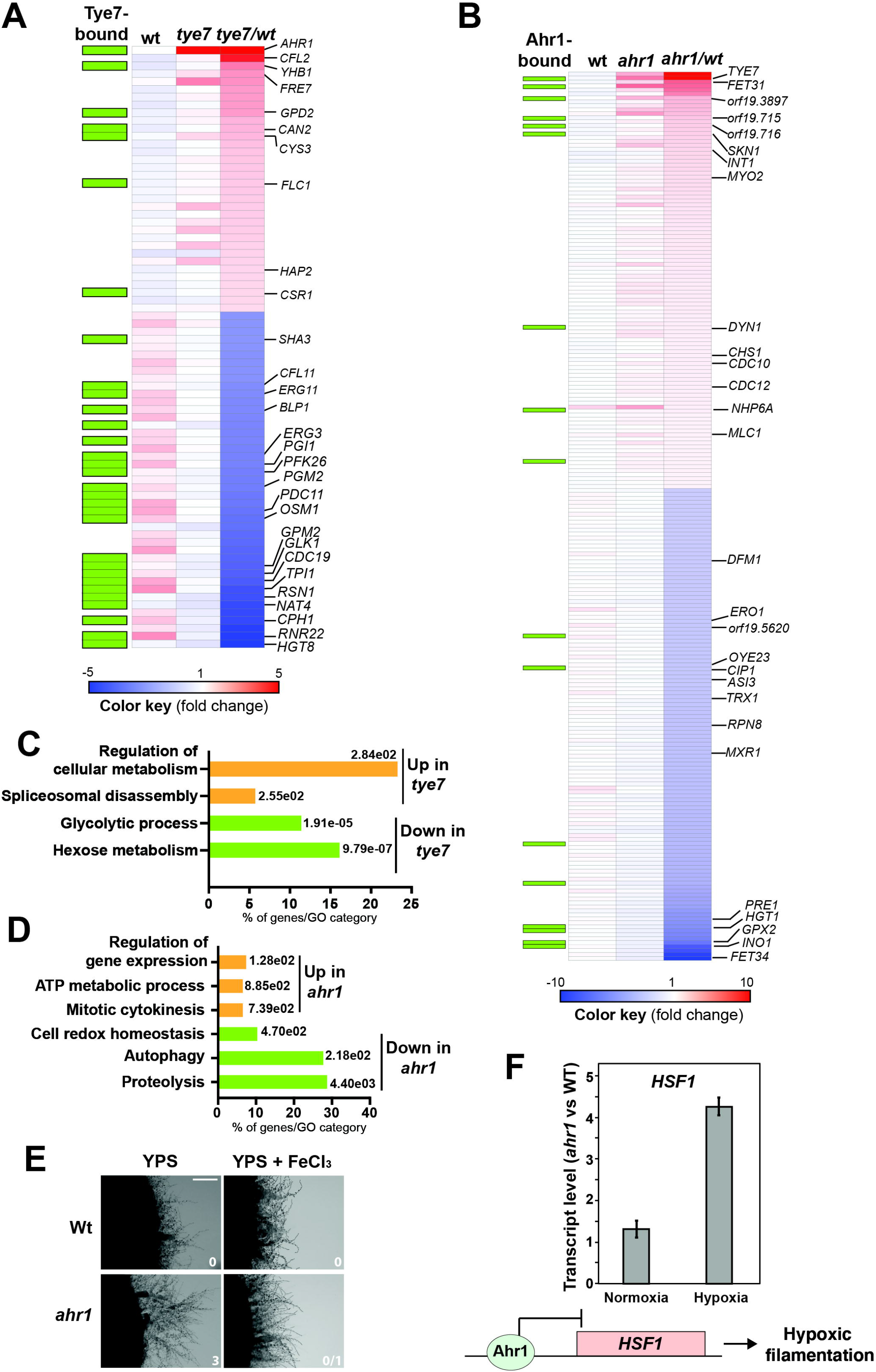
Tye7 and Ahr1 transcription factors act as a negative regulators of *C. albicans* hypoxic filamentation. Heatmap of differentially expressed transcripts in *tye7* (**A**) and *ahr1* (**B**) mutants. Plotted are all transcripts that were significantly differentially expressed (1.5-fold change cut-off and a 5% FDR) in *tye7* and *ahr1* mutants. Tye7- and Ahr1-dependant transcripts were identified by comparing the transcriptional profile of *tye7* or *ahr1* cells to that of WT cells. Relevant transcripts were annotated in the heatmap. The green boxes indicate gene promoters occupied by Tye7 and Ahr1 as previously defined by Askew *et al*. [26] and in the current study, respectively. Blue and red colors represent down- and up-regulated genes, respectively. (**C**,**D**) Gene function and biological process enriched in the transcriptional profiles of *tye7* (**C**) and *ahr1* (**D**). The *P*-value was calculated using a hypergeometric distribution with Multiple Hypothesis Correction, as described on the Gene Ontology (GO) Term Finder website. GO analysis was performed using the Candida Genome Database GO Term Finder. (**E**) Restauration of the WT hypoxic filamentation in *ahr1* mutant by supplementing YPS-agar medium with 100 µM FeCl_3_. Filamentation scores were indicated for each strain and tested condition. Bar, 20 µm. (**F**) Relative expression of *HSF1* in *ahr1* mutant. Transcript levels of *HSF1* were evaluated in *ahr1* cells exposed to normoxia and hypoxia relative to WT *HSF1* levels. Fold-inductions were calculated using the comparative CT method. A model of Ahr1 regulation of hypoxic filamentation through its repressive activity on *HSF1* promoter is shown.

The 34 transcripts upregulated in *tye7* mutant were enriched in iron utilization genes including the two ferric reductases *FRE7* and *CFL2*, the transcription factor *HAP2* and the heme transporter *FLC1* (**Figure 3A**). Tye7 has no apparent role in iron metabolism as it grew normally under iron-depleted media irrespective of oxygen abundance (**Supplementary Figure 1**). The activation of iron uptake transcripts was previously shown to accompany the yeast-to-hyphae transition and might reflect a phenomenon called adaptive prediction or predictive behaviour [40–43]. In *C. albicans*, such a concept implies a coactivation of process leading to filamentation together with other factors that are required during or after tissue invasion such as iron acquisition. Thus, activation of iron utilization transcripts in *tye7* might elucidate such adaptive anticipatory response as this mutant is hyperfilamentous.

### Transcriptional profiling of *ahr1* mutant under hypoxia uncover an iron starvation situation

Transcripts downregulated in *ahr1* mutant were enriched mainly in proteolysis (*ASI3, DFM1, PRE1, RPN8*) (**Figure 3B and 3D; Table S4**). This pattern might explain the hyperfilamentation phenotype of *ahr1* since genetic inactivation or pharmacological inhibition of the *C. albicans* proteosome were previously shown to induce filamentation in the absence of an inducing cue [44–49]. Repressed transcripts in *ahr1* also included genes related to oxidative stress such as the thioredoxin Trx1, the thiol reductase Ero1 and different proteins with oxidoreductase functions (Cip1, Mxr1, Oye23). Upregulated genes were enriched in function related to mitotic cytokinesis, ATP generation and gene expression regulation (**Figure 3D**). Activation of cytokinesis transcripts such as those encoding components of the septin ring (Cdc10, Cdc12, Chs1, Int1) and other actomyosin related proteins (Mlc1, Myo2, Dyn1) might mirror the enhanced hyphal growth of *ahr1*.

Transcript of the multicopper ferroxidase Fet34, a protein that oxidizes Fe^2+^ to Fe^3+^ for subsequent cellular uptake by transmembrane permease Ftr1 was highly repressed in *ahr1* (**Figure 3B**). Meanwhile, transcript of Fet31, another multicopper ferroxidase, was induced (**Figure 3B**) suggesting that Fet31 might be solicited to compensate for the repression of Fet34. While Fet31 was not dispensable for iron uptake in *C. albicans*, Fet34 plays an essential role in iron acquisition [50]. Thus, downregulation of Fet34 might reflect an impairment of iron homeostasis in *ahr1* which could also enable the observed hyperfilementation as previous work showed that iron starvation promotes hyphal growth [51]. To test this hypothesis, we first assessed *ahr1* growth in the presence of the iron-chelating agent BPS. Under yeast-promoting growth and either under hypoxia or normoxia, *ahr1* did not show any perceptible growth defect in the presence of BPS (**Supplementary Figure 1**). However, *ahr1* hyperfilamentation was reverted to a state comparable to that of the WT strain by supplementing the growth medium with 100 µM ferric chloride. This reinforces the hypothesis that the *ahr1* enhanced hyphal phenotype might reflect a nutritional scavenging response as a consequence of a depleted intracellular iron pool (**Figure 3E**).

To assess whether the differentially modulated transcripts in *ahr1* mutant are direct targets of this transcription factor, Ahr1 occupancy was assessed by ChIP coupled to high-density tiling arrays under hypoxic conditions. Only a few *ahr1*-misregulated transcripts have their promoter bound by Ahr1 (17/225 differentially expressed genes in *ahr1*) (**Figure 3B and Table S5**). This suggests that the observed *ahr1* hyperfilamentation phenotype is unlikely the result of expression alteration of Ahr1 direct targets. For instance, nor Fet34 or proteolysis gene promoters were bound by Ahr1. However, we found that Ahr1 bound the promoter of the heat shock transcription factor Hsf1, a transcriptional modulator of the Hsp90 chaperone network that is a master regulator of morphogenesis in *C. albicans* [52,53]. Former works by the Cowen group uncovered an intriguing phenomenon where either increasing or decreasing *HSF1* dosage promote filamentation through two independent mechanisms [54]. *HSF1* overexpression led to transcriptional activation of positive regulators of filamentation, including Brg1 and Ume6 while its depletion compromises Hsp90 function resulting in an increased filamentation. Since our microarrays analysis did not show noticeable alteration of *HSF1* expression, we used qPCR to assess the transcript level of *HSF1* in both WT and *ahr1* mutant strains. Under normoxia, *HSF1* was not modulated in *ahr1* as compared to the WT strain while under hypoxia the transcript level of *HSF1* was significantly increased in *ahr1* (**Figure 3F**). This suggest that, under hypoxia, Ahr1 act as a repressor on the promoter of *HSF1* which might promote hypoxic filamentation in *C. albicans*. Thus, Ahr1’s role as a repressor of hypoxic filamentation could be mediated through Hsf1 regulatory axis. These observations pave the road for future mechanistic studies investigating the high hierarchical role of Ahr1 as a transcriptional regulator of Hsf1-Hsp90 activity to control hypoxic filamentation. Intriguingly, our omics analysis supports a negative feedback loop control of Ahr1 and Tey7 as they bind the promoter of each other and were required to transcriptionally repress each other (**Figure 3A and 3B**).

### Genetic connectivity of the hypoxic filamentation network

Genetic interactions were used to assess functional relationships between the newly identified modulators of hypoxic filamentation including Ahr1, Tye7 and Sch9. Our data showed that the double mutant *ahr1tye7* had an additive phenotype as compared to their congenic strains suggesting that both Ahr1 and Tye7 act in parallel pathways (**Figure 4A**). Deleting *SCH9* in either *tye7* or *ahr1* resulted in slow-growing smaller colonies which hindered the assessment of filamentation score.

**Figure 4.**
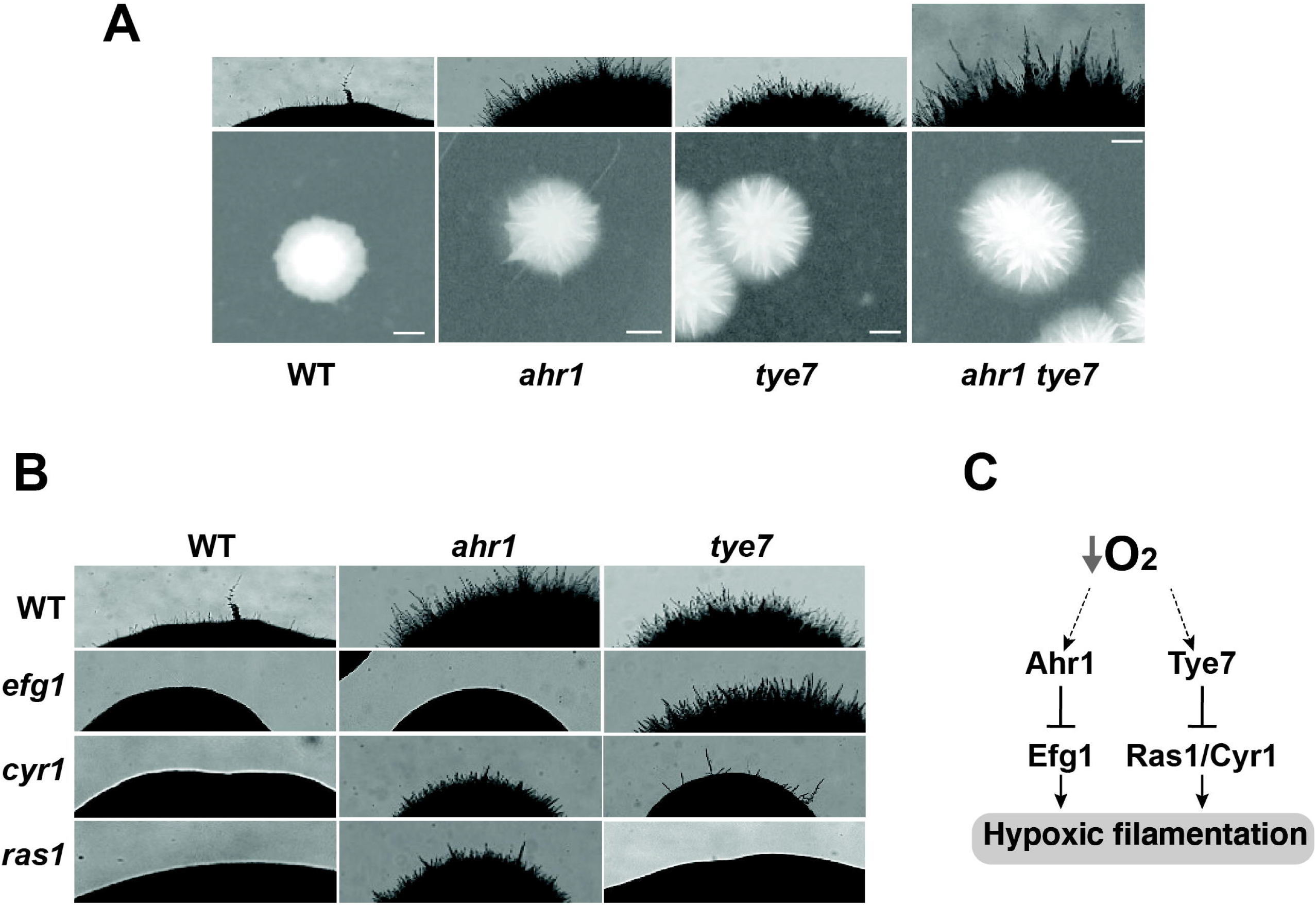
Genetic epistasis of the hypoxic filamentation circuit. (**A**) Ahr1 and Tye7 act in parallel pathways to modulate hypoxic filamentation. (**B**) Ahr1 and Tye7 repress the hypoxic filamentous growth via Efg1 and -cAMP-Ras1/Cyr1 pathways, respectively. *C. albicans* colonies were grown on a YPS-agar medium at 37°C under hypoxia (1% O_2_) for 3 days. (**C**) Schematic representation of the regulatory network that governs hypoxic filamentation in *C. albicans*.

The *C. albicans* morphogenetic switch is controlled by intertwined regulatory circuits that signal different filamentation cues. We tested the genetic interaction of Ahr1 and Tye7 with the high hierarchical regulators Efg1 and the Ras1/Cyr1 cAMP pathway known to promote filamentation in response to a myriad of stimuli [10]. Genetic inactivation of either Ras1 or Cyr1 in *ahr1* mutant has no apparent effect while deletion of *EFG1* completely abolished the hyperfilamentous phenotype of *ahr1* (**Figure 4B**). Contrary, the enhanced hyphal growth of *tye7* mutant under hypoxia was suppressed by either deleting *RAS1* or *CYR1* but not *EFG1* (**Figure 4B**). Taken together, these data suggest that Ahr1 and Tye7 repress the hypoxic filamentation growth via the Efg1 and Ras1/Cyr1 pathways, respectively. This finding also suggests that both Ahr1 and Tye7 might link the oxygen status of a colonized niche to the general core modulator of *C. albicans* filamentation such as Efg1 and Ras1/Cyr1 cAMP pathways.

In conclusion, the current study uncovered two new regulatory circuits that govern filamentation in response to oxygen levels. Both Tye7 and Ahr1 act as negative regulators of hypoxic filamentation and operate in two independent pathways as supported by our genetic interaction analysis and omics approaches.

## Materials and methods

### Strains, mutant collections, and growth conditions

The *C. albicans* strains used in this study were listed in **Table S6**. The kinase [24] and the transcriptional factor [23] mutant collections used for the genetic screens were acquired from the genetic stock center (http://www.fgsc.net). The transcriptional regulator [25] mutant collection was kindly provided by Dr. Dominique Sanglard (University of Lausanne). For general propagation and maintenance conditions, the strains were cultured at 30°C in a yeast-peptone-dextrose (YPD) medium supplemented with uridine (2% Bacto-peptone, 1% yeast extract, 2% dextrose, and 50 µg/ml uridine). Cell growth and genetic transformation were carried out using standard yeast procedures [56].

For gene expression profiling under hyphae-promoting conditions, cells were collected directly from agar plates with a cell scraper after growing for 48 h at 37°C under either normoxia (21% O_2_) or hypoxia (1% O_2_). Growth under hypoxic conditions was achieved by incubating agar plates in an anaerobic chamber (Oxoid; HP0011A) continuously flushed with nitrogen to set oxygen levels at 1% and to remove any gaseous by-products. Collected cells were rapidly frozen in liquid nitrogen and immediately processed for RNA extraction. The effect of iron chelation with BPS (Batho-phenanthroline disulfonic acid) on *C. albicans* growth was performed using spot dilution as described by Khemiri *et al*. [57].

### Genetic screen

For each mutant collection, strains were arrayed using a sterilized 96-well pin tool on Nunc Omni Trays containing YPS-agar and colonies were grown for four days at 37°C under normoxic (21% O_2_) and hypoxic conditions (1% O_2_). Plates were then imaged using the SP-imager system (S&P Robotics Inc.). A growth score was given for each mutant (**Table S1**). Each mutant hit was confirmed individually by assessing filamentation of at least five single cell-derived colonies.

### Expression Analysis by Microarrays and quantitative-RTPCR

Total RNA was extracted using an RNAeasy purification kit (Qiagen) and glass bead lysis in a Biospec Mini 24 bead-beater as previously described [58]. RNA was assessed for integrity on an Agilent 2100 Bioanalyzer prior to cDNA labeling. cDNA labeling and microarray procedure were performed as previously described [17]. Briefly, 20 µg of total RNA was reverse transcribed using 9 ng of oligo(dT)_21_ and 15 ng of random octamers (Invitrogen) in the presence of Cy3 or Cy5-dCTP (Invitrogen) and 400 U of Superscript III reverse transcriptase (ThermoFisher). After cDNA synthesis, template RNA was degraded by adding 2.5 U RNase H (Promega,) and 1µg RNase A (Pharmacia) followed by incubation for 20 min at 37°C. The labeled cDNAs were purified with a QIAquick PCR purification kit (Qiagen). DNA microarrays were processed and analyzed as previously described [59]. The GSEA PreRanked tool (http://www.broadinstitute.org/gsea/) was used to determine the statistical significance of correlations between the transcriptome of *C. albicans* hyphal cells under hypoxia with a ranked gene list as previously described [21,32].

For the *HSF1* qPCR experiment, a total of two biological and three assay replicates were performed. cDNA was synthesized from 60 ng of total RNA using High-Capacity cDNA Reverse Transcription kit (Applied Biosystems). The mixture was incubated at 25°C for 10 min, 37°C for 120 min and 85°C for 5 min. Two units per microliter of RNAse H (NEB) was added to remove RNA and samples were incubated at 37°C for 20 min. qPCR was performed using StepOne™ Real-Time PCR System (Applied Biosystems) and the PowerUp SYBR Green master mix (Applied Biosystems). The reactions were incubated at 95°C for 10 min and cycled for 40 times at 95°C, 15s; 60°C, 1 min. Fold-enrichments of each tested transcripts were assessed using the Ct comparative method and actin as a reference gene.

### Whole-genome location profiling by ChIP-chip

Ahr1 was TAP-tagged *in vivo* with a TAP-HIS1 PCR cassette in SN148 strain as previously described [60]. ChIP-chip of Ahr1 under hypoxic conditions was performed using tiling arrays as we have previously fulfilled [59]. Briefly, cells were grown as described for the microarrays experiment in YPS-agar plates, harvested with a cell scraper in microcentrifuge tubes and incubated for 20 min with 1% formaldehyde for DNA-protein crosslinking. Tiling arrays were co-hybridized with tagged immunoprecipitated (Cyanine 5-labeled) and mock immunoprecipitated (untagged SN148 strain; Cyanine 3-labeled) DNA samples. The hybridization was carried out at 42°C for 20 h in a Slide Booster Hyb-chamber SB800 (Advalytix), with regular micro-agitation of the samples. Slides were washed and air-dried before being scanned using a ScanArray Lite microarray scanner (PerkinElmer). Fluorescence intensities were quantified using ImaGene software (BioDiscovery, Inc.), background corrected and normalized for signal intensity (using Lowess normalization). The significance cut-off was determined using the distribution of log-ratios for each factor which was set at two standard deviations from the mean of log-transformed enrichments. Peak detection was performed using Gaussian edge detection applied to the smoothed probe signal curve, as described by Tuch et al. [61]. Both raw and processed microarray and ChIP-chip data have been submitted to the ArrayExpress database at EMBL-EBI (https://www.ebi.ac.uk/arrayexpress/) [62] under accession number E-MTAB-10882 and E-MTAB-10883, respectively.

## Abbreviations

ChIP-chip: Chromatin Immuno-Precipitation on chip
WT: Wild Type
YPD: Yeast-Peptone-Dextrose
YPS: Yeast-Peptone-Sucrose
cAMP: 3’,5’-cyclic AMP
GSEA: Gene Set Enrichment Analysis
GO: Gene Ontology
BPS: Batho-Phenanthroline diSulfonic acid
TAP: Tandem Affinity Purification
Ct: Cycle threshold.

## Acknowledgements

Not applicable.

## Authors’ contributions

Conceptualization: AS; Methodology: MH, AB, FT and AS; Funding acquisition: AS; Resources: AS; Supervision: AS and FT; Writing original draft: AS; Writing, review & editing: AS, MH, FT and AB. All authors listed gave final approval for publication.

## Availability of data and materials

Gene expression and ChIP-chip data are available at ArrayExpress database at EMBL-EBI (https://www.ebi.ac.uk/arrayexpress/) [62] under accession number E-MTAB-10882 and E-MTAB-10883.

## Ethics approval and consent to participate

Not applicable

## Consent for publication

Not applicable

## Supplementary data

**Supplementary Figure 1**. The *C. albicans* WT, *tye7* and *ahr1* strains were serially diluted, spotted on YPD-agar medium with different concentrations of BPS and incubated for 2 days at 30°C under normoxic (21% O_2_) or hypoxic (1% O_2_) environments.

**Table S1**. Raw data of the genetic survey for regulatory proteins required for hypoxic and normoxic filamentation

**Table S2**. Transcripts differentially expressed in *C. albicans* WT (SN250) colonies under hypoxia using a 1.5-fold change cut-off and a 5% false discovery rate.

**Table S3**. Gene Set Enrichment Analysis (GSEA) of the transcriptome of *C. albicans* WT (SN250) colonies under hypoxia.

**Table S4**. Transcripts differentially expressed in *C. albicans* in *tye7* and *ahr1* colonies under hypoxia using a 1.5-fold change cut-off and a 5% false discovery rate.

**Table S5**. List of Ahr1-bound intergenic regions as defined by ChIP-tiling arrays

**Table S6**. Lists of *C. albicans* strains and primers used in this study.

