## Supplementary figures and images for "Transcriptional control of hypoxic hyphal growth in the fungal pathogen *Candida albicans*"

### Supplementary Figure 1

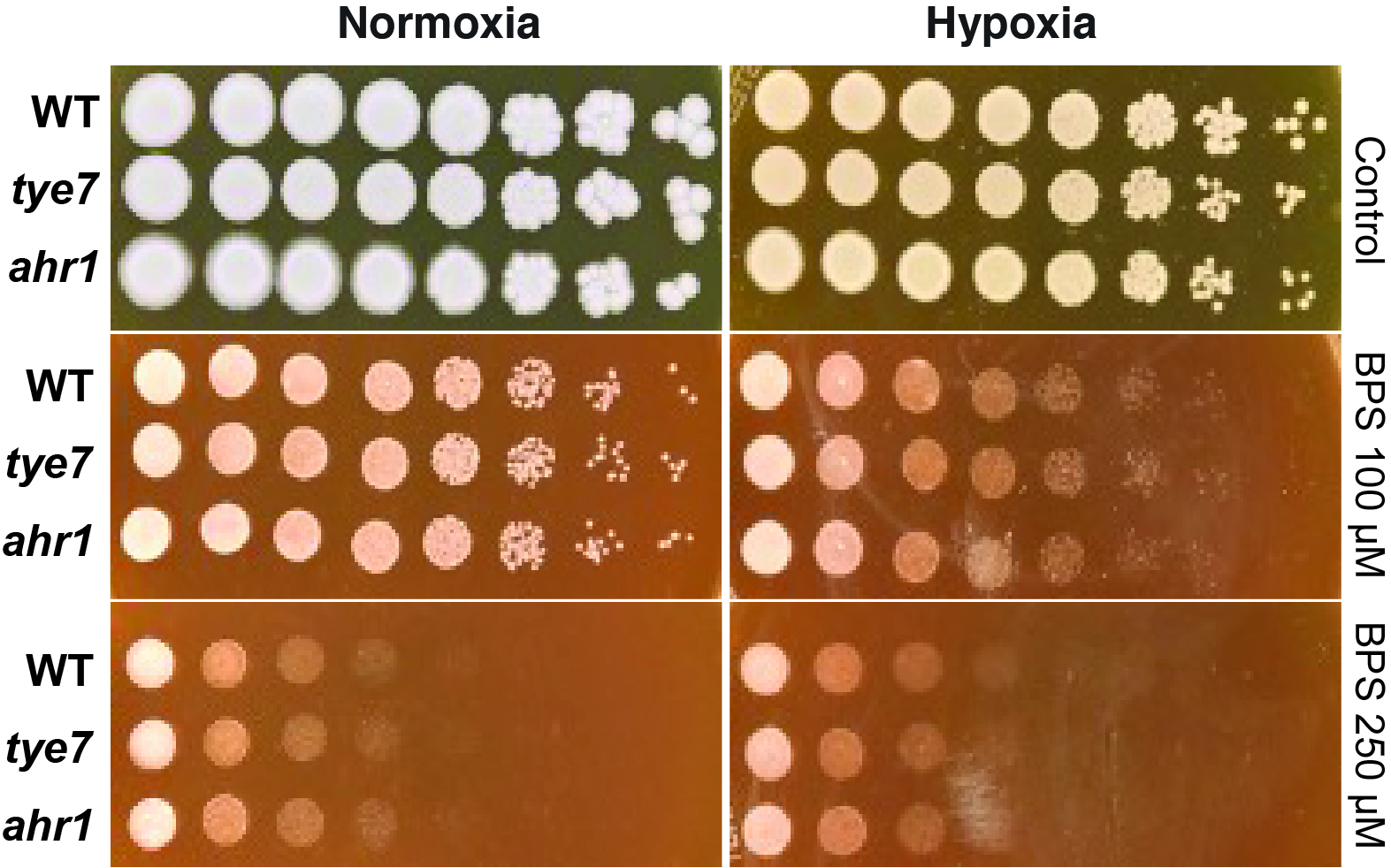
